# Site-Specific Noninvasive Delivery of Retrograde Viral Vectors to the Brain

**DOI:** 10.1101/2025.04.24.650522

**Authors:** Manwal Harb, Shirin Nouraein, Jerzy O. Szablowski

## Abstract

Neuronal activity underlies the brain function. Different behaviors, physiological processes, and disorders depend on which neurons are active at a given moment. Treating brain disorders without side effects will require exclusive control of disease-relevant neurons. Traditionally, small molecule drugs could control a subset of neurons that express a molecularly specific receptor. Local noninvasive therapies such as delivery of neuromodulatory agents with focused ultrasound blood-brain barrier opening (FUS-BBBO) also added spatial precision allowing one to control specific brain regions without surgery. However, the final characteristic of neurons, which other neurons they connect to, remains underexplored as a therapeutic target. If targeting neurons based on their connectivity was possible noninvasively, it would open the doors to broadly deployable precise therapies that can target selected subgroups of neurons within a brain region. Such delivery could be achieved with retrograde-tracing adeno-associated viral vectors (AAVs). For noninvasive delivery with FUS-BBBO, AAV9 has emerged as the most promising serotype. However, its retrograde-tracing version, the AAV9.retro, has not been evaluated for FUS-BBBO delivery. Here, we show that following such noninvasive delivery AAV9.retro can safely transduce neuronal projections with comparable efficiency to a direct intracranial injection. Compared to AAV8, a naturally occurring vector with low retrograde transduction, AAV9.retro offers superior retrograde transduction and comparable transduction at the site of delivery. Overall, we show that AAV9.retro is a valuable FUS-BBBO gene delivery vector, while also highlighting the surprising possibility of improved specificity of transduction of projections compared to invasive delivery.

## INTRODUCTION

Brain is interconnected through groups of neurons that extend their axons into distal sites. These connections, or projections, integrate information from multiple brain regions and perform basic brain functions such as cognition, motor control, or memory^1-3^. Targeting projections can control highly specialized behaviors, for example underlying various aspects of depression^4^, reward-seeking^5^, or pain sensation^6^.

Recent discovery of viral vectors that can be transported through axonal projections opened important areas of investigation and a highlighted their potential for brain therapy^7-9^. Two major classes of such vectors exist. Vectors targeting anterograde projections^10,11^ enter through the dendrites, are transported through axons, and diffuse across the synapse to transduce the next neuron in the pathway. Retrograde-tracing vectors, such as AAV2.retro^7,12^ or AAV9.retro^13^, can be uptaken within the axon and then transported to the cell body and nucleus to express the genetic cargo. In consequence, with a retrograde-targeting vector, one can target a brain site, and transduce cell bodies within distant regions that project to the targeted site.

Retrograde vectors, administered invasively through a direct intraparenchymal (**IP**) injection, catalyzed several important discoveries in neuroscience. For example, projections from the prefrontal cortex to the nucleus accumbens (**NAcc**) showed acute avoidance behavior when the NAcc was inhibited^14^, while projections from the ventral tegmental area (**VTA**) to the basolateral amygdala controlled anxiety-like behaviors^15^. These discoveries raised the potential for development of gene therapies that could resolve many brain disorders that are based on the abnormal regulation of the brain circuitry, rather than specific cell types, or brain sites. Such control could be achieved, for example, through chemogenetic regulation^16^ or by resetting their excitability from a pathogenic to a well-controlled state through expression of inhibitory ion channels^17^. However, delivery of these viral vectors to the brain remains a challenge. Since the neuronal pathways project to specific brain sites, the delivery must be spatially specific. The common methods of gene delivery, such as direct intraparenchymal injections, require a complex brain surgery, risk side effects such as infection or hemorrhage, and require patients who are willing and able to undergo surgery^18^. On the other hand, new vectors that can transcytose from the blood into the brain across the blood-brain barrier (**BBB**) noninvasively, but currently lack spatial precision, preventing site-specific transduction of spatially defined brain circuits. Over the past two decades, Focused ultrasound BBB opening (**FUS-BBBO**) emerged as a promising solution for targeting neuronal circuits. FUS-BBBO enables non-surgical, systemic delivery of AAVs from the blood into the targeted brain sites with millimeter precision. FUS-BBBO was also shown to be safe and effective in various species including mice^16^, rats^19^, rabbits^20^, non-human primates^21^ and humans^22^. In each case, FUS-BBBO could deliver various cargo such as small molecules^23^, proteins^24^, AAVs^16^ and nanoparticles^25^.

Recent studies have shown that AAV9 has higher transduction efficiency compared to other AAV serotypes, including AAV2 and its retrograde tracing variant AAV2.rg^12^. However, to date, the efficiency of the retrograde tracing variant of AAV9 with FUS-BBBO delivery has not been tested.

Here, we evaluate the transduction efficiency of a retrograde-tracing vector, AAV9.retro^13^, delivered to the brain through FUS-BBBO, comparing its transduction efficiency at the site of delivery and projections to the invasive IP injection. Both modes of delivery used a non-tracing AAV8 benchmark. We found improved retrograde tracing for AAV9.retro compared to the AAV8 control with either mode of delivery. We also found that FUS-BBBO transduction of neuronal pathways with AAV9.retro resulted in no neuronal cell loss and has not increased the presence of markers of inflammation. Interestingly, in one of the brain regions, NAcc, the transduction of the cell bodies at the site of delivery was lower for FUS-BBBO, while maintaining comparable transduction of projections. This result suggests FUS-BBBO improved specificity of retrograde tracing compared to IP injection. Overall, this study shows successful transduction of retrograde projections by AAV9.retro with efficiency that was comparable between IP and FUS-BBBO deliveries, and superior at retrograde tracing compared to AAV8 in both modes of delivery.

## RESULTS

To evaluate the performance of our retrograde-tracing vector following noninvasive delivery, we selected regions of the brain that enabled identification of retrograde transduction in previous studies^12^. Specifically, we selected the nucleus accumbens (NAcc) and the basal pontine nucleus (BPN) as two deep-brain regions which are projected to from the cortical and olfactory areas (**Fig. 1**). We used AAV8 as a control vector with low retrograde tracing ability^7^ both to confirm the spatial accuracy of FUS-BBBO delivery and to provide benchmark of retrograde tracing ability for AAV9.retro.

**Figure 1.**
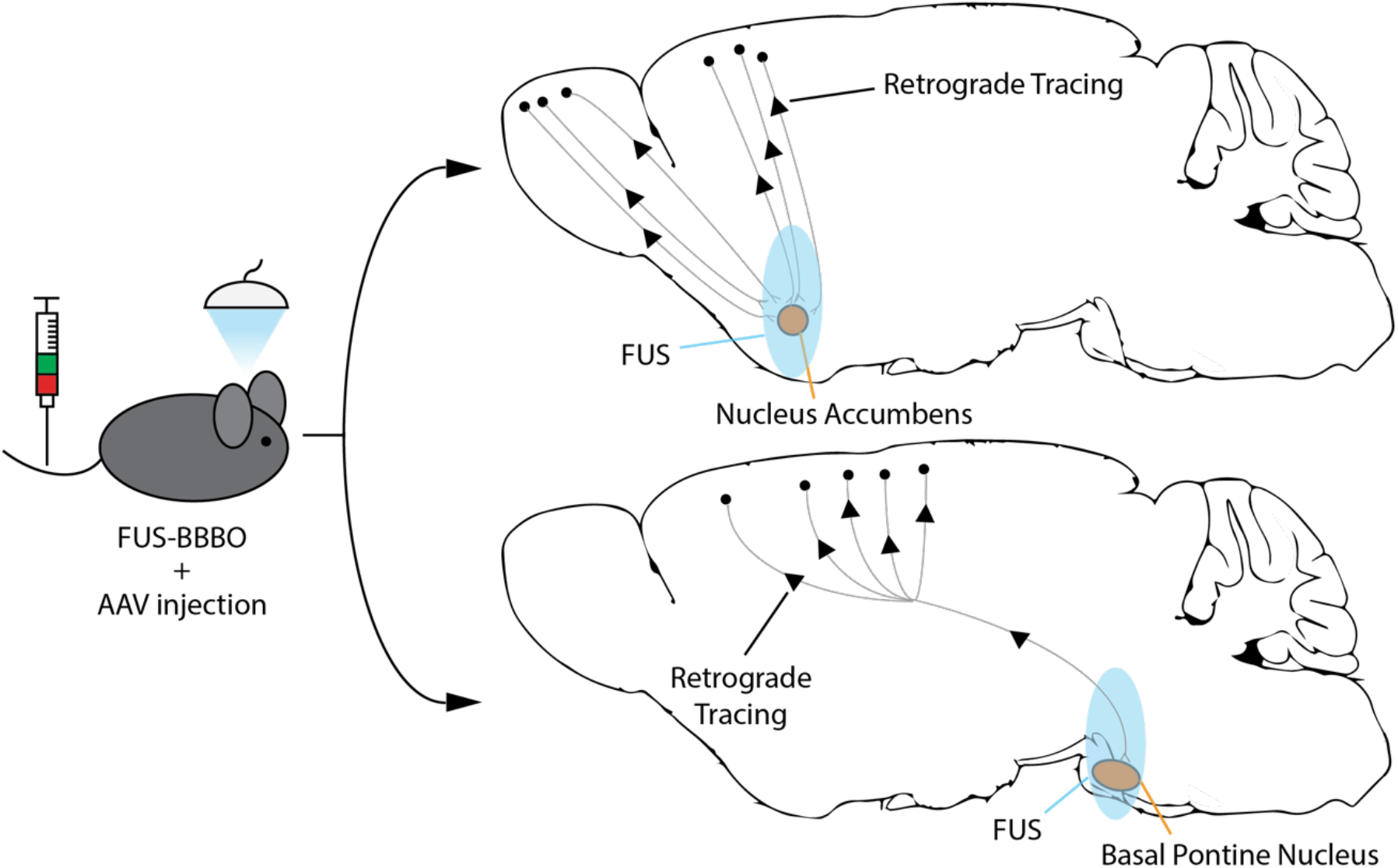
Delivery of Retrograde-Tracing AAVs following Focused ultrasound blood-brain barrier opening (FUS-BBBO). The ultrasound-enhanced opening of the BBB allows the systemically administered AAVs to enter from the circulation into the targeted brain site. By targeting deep-brain areas such as the nucleus accumbens (NAcc) and the basal pontine nucleus (BPN), AAVs with retrograde-tracing capability can be delivered and transduce neurons in spatially distinct brain regions such as those on the brain’s surface (cortex and olfactory bulb).

### Transduction of the retrograde projections of the nucleus accumbens

To evaluate the efficiency of transduction AAV9.retro and AAV8, we first performed FUS-BBBO at the NAcc of wild-type mice to deliver AAVs. We placed EGFP and mCherry under the neuron-specific human Synapsin (hSyn) promoter and packaged each plasmid into AAV9.retro (hSyn-EGFP) and AAV8 (hSyn-mCherry). We delivered the AAVs to the brain using ultrasound parameters based on previous studies^27^ and experimental optimization for our FUS system. In a pilot experiment we found that 1.2 MPa peak pressure at 1.5 MHz frequency, with 1 Hz pulse repetition frequency for 120 pulses resulted in successful BBB opening (**Supplementary Figure S1**). Immediately following FUS-BBBO, we injected 10^10^ viral particles per gram of body weight of each virus. Mice were euthanized 2 weeks later, with coronal brain sections taken (40 μm thickness). As a positive control, we co-delivered the AAVs through a direct intraparenchymal (IP) injection into the NAcc at a dose of 4×10^8^ viral particles per gram of body weight of each vector.

We observed successful gene delivery at the targeted FUS site (**Fig. 2 a,d**), with no significant difference in transduction efficiency between AAV8 (mean of 59.2 ± SEM of 27.69 mCherry-positive cells per section) and AAV9.retro (mean of 95.6 ± SEM of 29.97 EGFP-positive cells per section; paired two-tailed *t* test, n=5). Similarly, IP injection showed comparable transduction at the site of delivery as well, with a mean of 467.7 ± 145.8 mCherry-positive cells and 476.6 ± 83.88 EGFP-positive cells per section at the targeted site for AAV8 and AAV9.retro respectively (mean ± SEM, paired two-tailed *t* test, n=5). For both FUS-BBBO and IP delivery groups, we observed EGFP expression in spatially distinct areas of the brain that have neurons projecting into the NAcc, namely the cortex (**Fig. 2b,e**) and the olfactory area (**Fig. 2c,f**). In the FUS group, we observed more AAV9.retro transduction with 57.92 ± 19.35 EGFP-positive cells in the cortex on average compared to 8.94 ± 2.5 mCherry-positive cells (6.482-fold difference, p = 0.0493, paired two-tailed *t* test, n=5). The IP group showed 46.5 ± 11.49 EGFP-positive cells in the cortex compared to 2.06 ± 1.12 mCherry-positive cells (18.6-fold difference, p = 0.0085 paired two-tailed *t* test, n=6). Similarly, in the olfactory area, 176.8 ± 49.64 cells were transduced by AAV9.retro following FUS-BBBO compared to 13 ± 7.45 cells transduced by AAV8 (13.6-fold, p = 0.0378, paired two-tailed *t* test, n=4). Meanwhile, when delivered by IP injection, AAV9.retro transduced on average 152.8 ± 31.57 cells, which is significantly higher than the 0.75 ± 0.48 cells transduced by AAV8 on average (203.73-fold, p = 0.0175, paired two-tailed *t* test, n=4). When comparing FUS-BBBO delivery of AAV9.retro relative to IP injection, there were significantly fewer transduced cells at the site of delivery (**Fig. 2g**), with 0.233-fold difference observed (p = 0.0048, unpaired two-tailed *t* test with Welch’s correction, n=5 for FUS group and n=6 for IP group). AAV9.retro transduction in the cortex (**Fig. 2h**) and olfactory area (**Fig. 2i**), however, did not differ significantly between the two delivery routes, where AAV9.retro transduction was 1.246-fold and 1.157-fold following FUS-BBBO respectively relative to IP injections (unpaired two-tailed *t* test, n=5 for cortex FUS group, n=6 for cortex IP group and n=4 for both olfactory area groups). This translates to a significant improvement in specificity of transduction of projections, as defined by the ratio of the numbers of transduced cells at the projection sites and at the site of AAV delivery. We found that AAV9.retro improves the specificity of projection transduction in the cortex by 6.027-fold (p = 0.0219, ratio paired two-tailed *t* test, n=5 for FUS and n=6 for IP) and in the olfactory area by 6.093-fold (p = 0.0035, ratio paired two-tailed *t* test, n=4 for both groups) when FUS-BBBO delivery is performed compared to IP injection.

**Figure 2.**
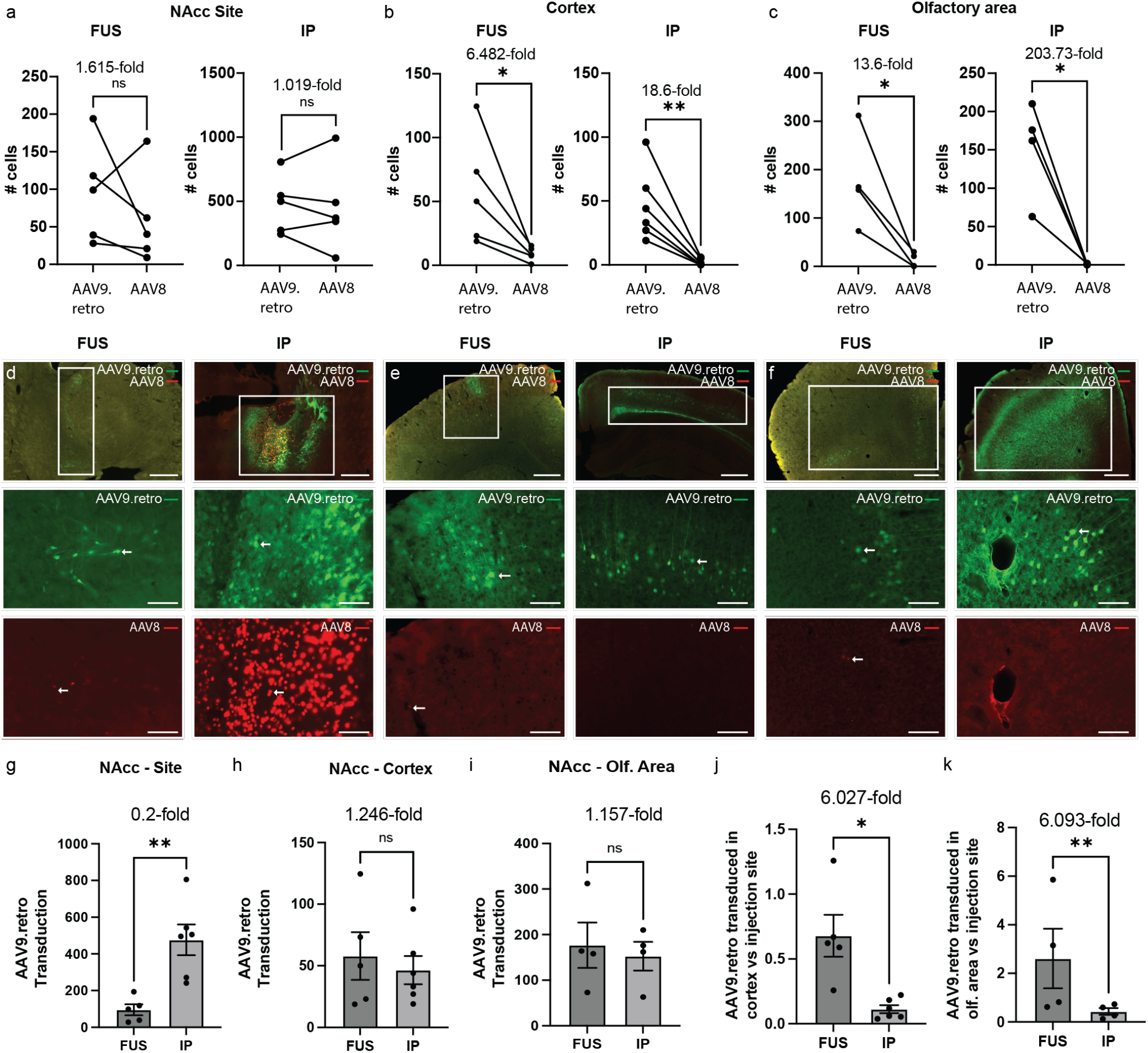
Retrograde transduction following targeting of NAcc. **a)** Comparable AAV9.retro and AAV8 expression was observed at the targeted NAcc site following FUS-BBBO (n=5, p = 0.3648, paired two-tailed t test) and following intraparenchymal injection (n=6, p = 0.9259, paired two-tailed t test). In both FUS and IP groups, AAV9.retro expression was significantly greater in the **b)** cortex (n=5, p = 0.0493 for FUS; n=6, p = 0.0085 for IP; paired two-tailed t test) and **c)** olfactory area (n=4, p = 0.0378 for FUS; n=4, p = 0.0175 for IP; paired two-tailed t test) compared to the AAV8 control. **d)** Representative images showing AAV9.retro (green) and AAV8 (red) transduction at the NAcc following FUS and IP injection. **e)** Transduction at cortex and **f)** olfactory area following gene delivery to NAcc. Arrows indicate examples of positively transduced cells. **g)** IP injections to NAcc showed significantly higher AAV9.retro transduction compared to FUS-BBBO delivery at the targeted site (n=5 for FUS and n=6 for IP; p = 0.0048, unpaired two-tailed t test with Welch’s correction). **h)** There were no significant differences in transduction between the delivery routes at the untargeted sites of the cortex (n=5 for FUS group, n=6 for IP group, p = 0.6098, unpaired two-tailed *t* test) or **i)** olfactory area (n=4 for both groups, p = 0.6975, unpaired two-tailed *t* test). **j)** The ratio of AAV9.retro transduction at the cortex relative to the site of delivery was significantly greater following FUS-BBBO compared to IP injection (n=5 for FUS and n=6 for IP, p = 0.0219, ratio paired two-tailed t test). **k)** AAV9.retro transduction was also significantly greater at the olfactory area compared to the NAcc when delivered by FUS-BBBO (n=4 for both groups, p = 0.0035, ratio paired two-tailed t test). All values are mean ± SEM. Scale bars are 500 μm for top row (overlay) panels and 100 μm for middle (AAV9.retro, green) and bottom row (AAV8, red) panels. (** = p<0.01, * = p<0.05, ns = not significant).

### Transduction of the retrograde projections of the basal pontine nucleus

We then performed FUS-BBBO at the BPN as an alternative site for targeting cortical neuron projections. The layer V pyramidal neurons in the cortex project to the BPN^28^, and can be visualized through sagittal sections. We observed successful gene delivery with no significant difference in transduction between the two vectors at the targeted sites for both FUS-BBBO and IP injection delivery (**Fig. 3a,c**). Per tissue section, FUS-BBBO-delivered AAV9.retro transduced 116.3 ± 65 and AAV8 123.8 ± 95.52 neurons (mean ± SEM, p = 0.6982, n=6, paired two-tailed *t* test). IP injection led to transduction of 1795 ± 711.2 neurons through AAV8, and 585 ± 240.7 neurons through AAV9.retro (mean ± SEM, p = 0.0687, n=6, paired two-tailed *t* test). For both delivery modes, EGFP expression, indicative of AAV9.retro transduction, was present in the cortex (**Fig. 3b,d**). In FUS-BBBO a mean of 408.5 ± 147 neurons were transduced per section. In the same group of mice, expression of mCherry, indicative of transduction by AAV8, was present in 1.83 ± 0.75 neurons per section (mean ± SEM., p = 0.0398, paired two-tailed *t* test, n=6). While for IP injection, 611.7 ± 188.1 EGFP-positive neurons were quantified on average in the cortex, there was no substantial mCherry-positive expression observed (mean ± SEM, p = 0.00227, paired two-tailed *t* test, n=6). FUS-BBBO delivery of AAV9.retro relative to IP injection showed no significant difference in transduced cells at the site of delivery (**Fig. 3e**), with 0.199-fold difference observed (unpaired two-tailed t test with Welch’s correction, n=6 for each group). Similarly, AAV9.retro transduction in the cortex (**Fig. 3f**) also showed no significant difference between the two delivery routes, where AAV9.retro transduction was 0.668-fold following FUS-BBBO relative to IP injections (unpaired two-tailed t test, n=6 each group). The results also show that the ratio of AAV9.retro transduction at the cortex relative to the BPN (**Fig. 3g**) does not differ significantly between the delivery routes (ratio paired two-tailed *t* test, n=6 for both groups), unlike when the NAcc was targeted.

**Figure 3.**
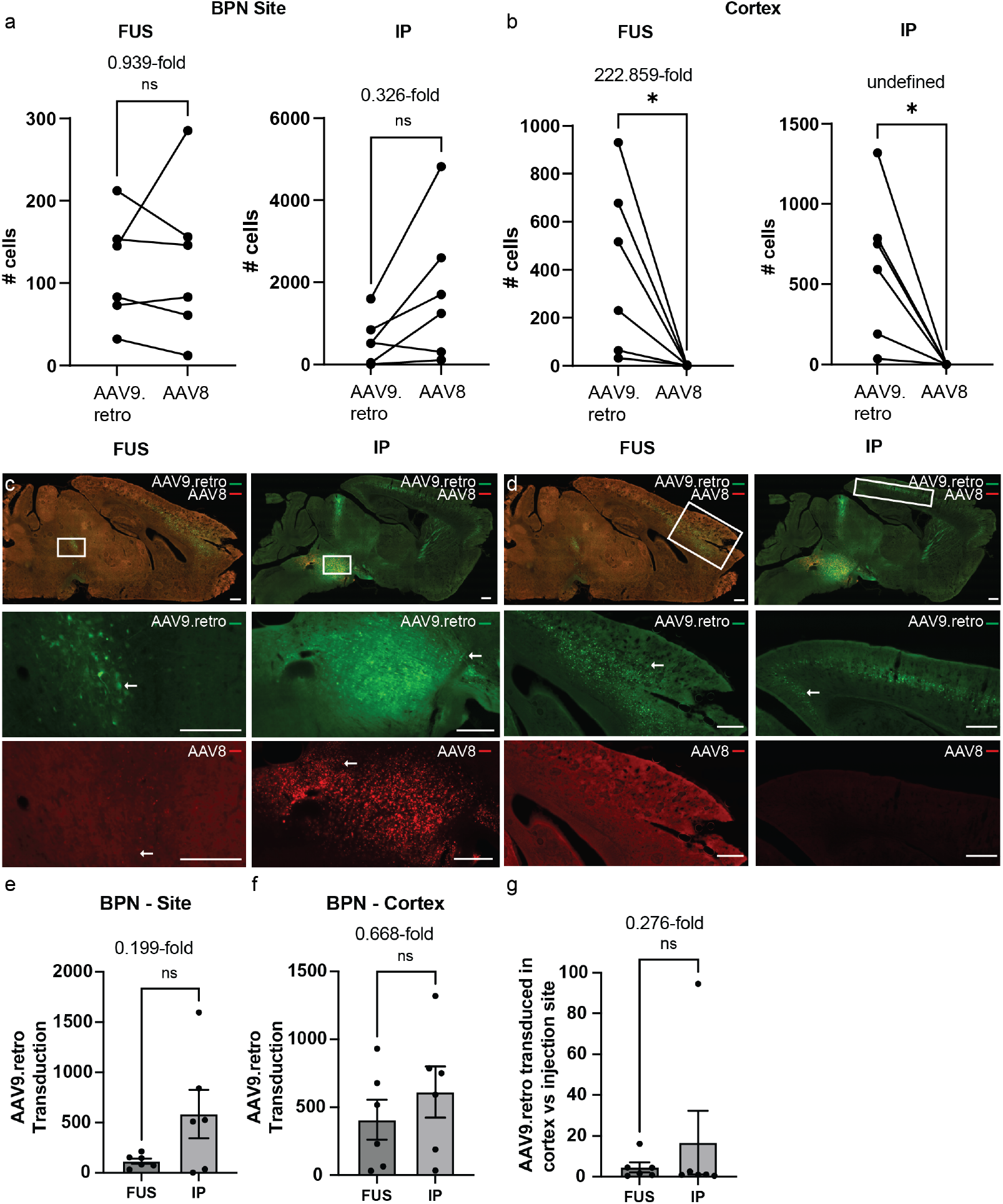
Retrograde transduction following FUS-BBBO targeting of BPN. **a)** AAV9.retro and AAV8 expression were observed at the targeted BPN site following FUS-BBBO (n=6, p = 0.6982, paired two-tailed t test) and following IP injection (n=6, p = 0.0687, paired two-tailed t test). **b)** AAV9.retro expression was significantly greater in the cortex for both FUS group (n=6, p = 0.0398, paired two-tailed t test) and the IP group (n=6, p = 0.0227, paired two-tailed t test) compared to the AAV8 control. Representative images showing AAV9.retro (green) and AAV8 (red) transduction at the **c)** targeted site (BPN) and **d)** cortex. **e)** The targeted BPN and **f)** the non-targeted cortex There were no significant differences in AAV9.retro transduction when delivered by either delivery method to the **e)** targeted BPN (n=6, p = 0.1094. unpaired two-tailed *t* test with Welch’s correction) and **f)** non-targeted cortex (n=6, p = 0.4147. unpaired two-tailed *t* test). **g)** The ratio of AAV9.retro transduction at the cortex compared to the BPN was not significantly different when performing FUS-BBBO or IP injection (n=6 for each group, p = 0.9595, ratio paired two-tailed t test). Arrows indicate examples of positively transduced cells. All values are mean ± SEM. Scale bars are 500 μm. (*p<0.05, ns = not significant).

### BBB permeability of AAV8 and AAV9.retro

We also compared AAV transduction at the site of delivery to the contralateral site to test whether the vectors were capable permeating through the intact BBB or diffusing away from the site of delivery. Since the numbers of positive cells in the contralateral non-targeted sites were often zero and not-normally distributed, we used an unpaired non-parametric Mann-Whitney test for all comparisons in this section. Following FUS-BBBO **(Fig. 4a-d)**, the targeted sites exhibited significantly higher transduction than the contralateral sites for all vectors and both in NAcc (p = 0.0079 for both comparisons, n=5) and BPN (p = 0.0022 for both comparisons, n=6). Similar observations were detected following IP injections **(Fig. 4e-h)** as there were significant transduction at the ipsilateral side relative to the contralateral side (p = 0.0022 for all comparisons, n=6 for each of NAcc and BPN groups).

**Figure 4.**
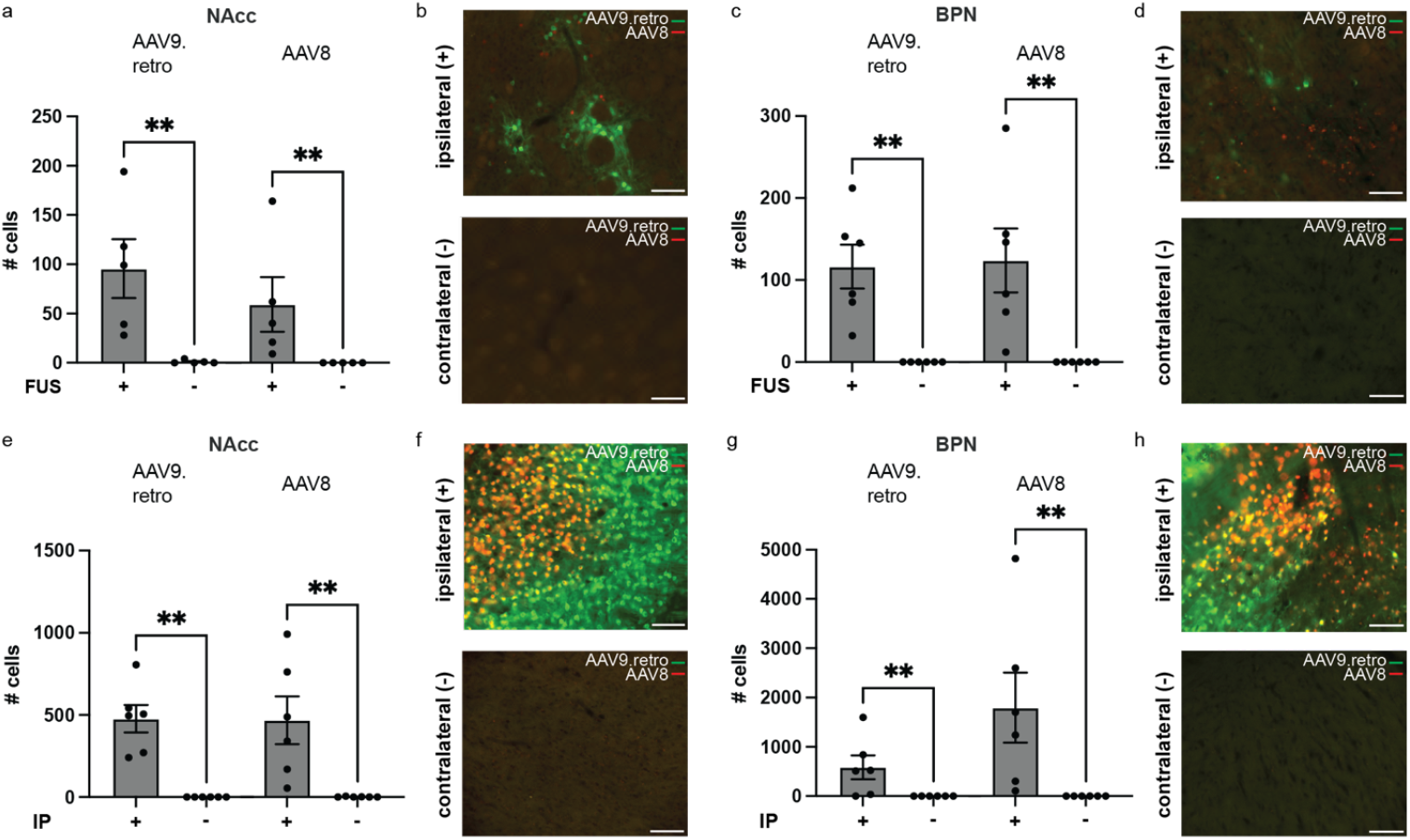
AAV9.retro and AAV8 are spatially restricted to the site of delivery. **a, b)** AAV9.retro and AAV8 expression were observed at the targeted NAcc site but not at the contralateral control following FUS-BBBO (n=5, p = 0.0079, unpaired two-tailed Mann Whitney test), suggesting no passage of either vector through an intact BBB. **c, d)** Similar results were observed for the BPN site targeting (n=6, p = 0.0022, unpaired two-tailed Mann Whitney test). Similarly, expression of both vectors was restricted to the site of IP injection at the **e, f)** NAcc (n=6, p = 0.0022 for both comparisons, unpaired two-tailed Mann Whitney test) and **g, h)** BPN (n=6, p = 0.0022 for both comparisons, unpaired two-tailed Mann Whitney test). **b, d, f, h)** Representative images showing transduction of AAV9.retro (green) and AAV8 (retro) following FUS-BBBO or IP injections at NAcc and BPN sites. All values are mean ± SEM. Scale bars are 100 μm. (**p<0.005).

### Safety and cellular toxicity of AAV9.retro and AAV8

To test the safety of the procedures and viral vectors, we documented the weight of the mice for up to 4 days following procedures. We observed no significant loss in weight (<10%) for both NAcc and BPN mice groups **(Fig. 5a,b)**. We also compared both the targeted and the contralateral sites to observe whether there was cellular loss at the site of delivery **(Fig. 5c-f)** as evaluated by the numbers of DAPI-positive cells. We found no significant differences in the number of cells between ipsilateral and contralateral sites following either route of delivery at both targeted regions (unpaired two-tailed *t* test, n=5 for NAcc group with FUS-BBBO, n=6 for all remaining groups).

**Figure 5.**
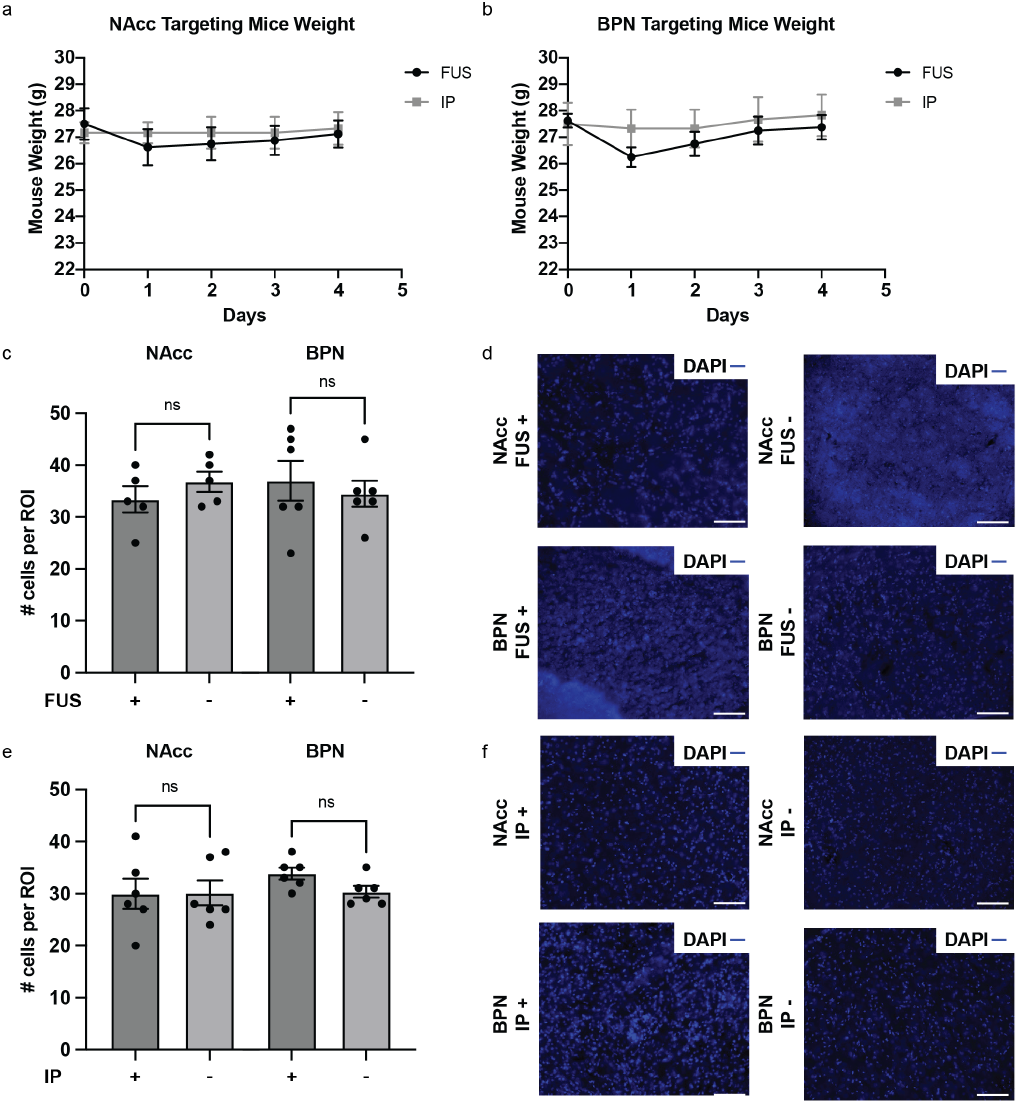
FUS-BBBO and IP delivery result in no substantial weight loss or cell loss at the targeted site. **a-b)** Mice did not experience significant weight loss (<10%) following either delivery route or AAV injections, making full recovery from the procedures. **c)** There were no significant differences in the number of DAPI-positive cells per region of interest (ROI; 100 μm x 100 μm bounded square at the site of delivery) between the targeted site and the contralateral control following FUS-BBBO at the NAcc (n=5, p = 0.3181, un-paired two-tailed t test) and at the BPN (n=6, p = 0.5983, un-paired two-tailed t test). **e)** IP injections of AAVs showed no significant difference in cellular loss per ROI compared to the contralateral control at both the NAcc (n=6, p = 0.9654, un-paired two-tailed t test) and the BPN (n=6, p = 0.0502, un-paired two-tailed t test). **d-f)** Representative images showing nuclear staining (DAPI, blue) of cells at the delivered sites and at the contralateral sites. All values are mean ± SEM. Scale bars are 100 μm. (ns = not significant).

## DISCUSSION

Retrograde-tracing viral vectors have shown promise in targeting specific neuronal circuits in the brain and modulating specific behaviors^1-5^. However, most of the gene delivery methods to the brain are either invasive or lack regional specificity^29-34^. FUS-BBBO has emerged as a safer noninvasive alternative for gene delivery to the brain, shown to target projections using vectors based on AAV2^12,16,27,35-38^. Compared to AAV2, the AAV9 serotype has exhibited improved transduction efficiency in the brain after FUS-BBBO and has undergone extensive study and engineering to further improve its transduction efficiency and spread, including a development of retrograde variant of its own^7,12,13,27,30^.

In this study, we illustrated the feasibility of using FUS-BBBO to deliver AAV9.retro to the brain and evaluate its transduction performance compared to a non-tracing control. The results highlight AAV9.retro’s capacity to safely transduce surface-level brain areas such as the cortex and olfactory bulb in a retrograde manner after being initially delivered by FUS-BBBO to deep brain regions such as the NAcc and the BPN. Moreover, while targeting either site resulted in cortical transduction, the neurons that project to these sites originate from different subpopulations, with layer II and V/VI neurons projecting to the NAcc and layer V neurons projecting to the BPN^28,39^. It is therefore possible to target specific cortical layers noninvasively with FUS-BBBO, which cannot be achieved by simply targeting FUS-BBBO to the cortex with non-tracing vectors. Approaches for cortical layer-specific targeting and modulation by targeting one indirect site in the brain could enable new treatments that are not currently possible. For example, layer V of the primary motor cortex is known to be involved in pain processing^40^. Targeting the cortical layer V neurons with therapeutics, for example with Acoustically Targeted Chemogenetics (ATAC)^16^, could enable new noninvasive therapies for pain management.

Our results revealed that IP injection resulted in significantly higher transduction at the targeted site with AAV9.retro compared to FUS-BBBO (5.1-fold for NAcc, and 5.0-Fold for BPN). However, the numbers of retrograde-transduced cells were not statistically significantly different between the delivery methods (1.3-fold for NAcc, and 0.7-fold for BPN, for AAV9.retro). Thus, despite lower AAV9.retro delivery to the site by FUS-BBBO, the retrograde transduction was comparable for IP and FUS-BBBO delivery in both cases. This raises an interesting prospect – FUS-BBBO delivery may be more specific for transduction of projections compared to the IP delivery. Further analysis showed the ratio of transduction of the projection sites compared to the site of interest was only statistically significant for NAcc, and not BPN. Further work will be needed to determine the mechanisms of this potential enhancement of projection transduction specificity.

FUS-BBBO delivery requires substantially higher doses of AAVs compared to a direct injection. In this study, IP injections used a total of AAV8 was 4×10^8^ viral genomes (VG), while intravenous dose of AAV9.retro was substantially higher – 1×10^10^ VG per gram of body weight of a mouse, or 2.5×10^11^ VG total for a 25-gram mouse. In our previous studies, we have shown more than 50% transduction efficiency at the site of FUS-BBBO at a tenfold lower dose when using an engineered vector, which we called AAV.FUS.3^27^. Thus, further engineering of AAV9.retro capsid could optimize transduction efficiency following FUS-BBBO reducing the overall dose needed and enhancing the transduction. For studies of large animals such as non-human primates, or in the clinic, improvement in transduction efficiency will reduce the high cost needed for the studies or any future therapies^41^.

AAV2.rg was shown to transduce the spleen and kidney at significantly higher levels than AAV2^12^. To alleviate potential confounds from peripheral expression, we used a neuron-specific synapsin promoter to concentrate expression only in the neuronal cells with our AAV9. This specificity was confirmed after finding no detectable transduction in other cell types in the brain such as astrocytes, oligodendrocytes and microglia/macrophages (**Supplementary Figure S2**) and in peripheral tissues such as the liver, kidney and spleen (**Supplementary Figure S3**. Future studies could employ a more ubiquitous promoter such as CAG to investigate AAV9.retro’s tropism to various peripheral tissues and cell types, which too could be optimized with AAV engineering.

## CONCLUSIONS

Overall, this study quantifies the transduction of a retrograde tracing AAV9.retro after FUS-BBBO delivery in mice in comparison to a naturally occurring AAV8. Surprisingly, we find that AAV9.retro showed comparable transduction of the retrograde projections for both intraparenchymal and FUS-BBBO delivery, despite significant differences in transduction at the site of original AAV delivery. Our findings also demonstrate that large swathes of the specific layers of the cortex could be targeted by noninvasively targeting a single deep brain site with FUS-BBBO.

## MATERIALS AND METHODS

### Animals

Male C57BL/6J mice between 10 and 14 weeks old were obtained from Jackson Laboratory (Bar Harbor, ME, USA). Animals were housed in a 12-hour light/dark cycle and were provided with water and food ad libitum. All experiments were conducted under a protocol approved by the Institutional Animal Care and Use Committee (IACUC) of Rice University.

### Plasmid and Viral Vectors

Plasmids were either obtained or modified from Addgene. The plasmid, pAAV-hSyn-EGFP (Addgene #50465) was packaged into an AAV9.retro vector^13^ using a commercial service (Baylor College of Medicine Gene Vector Core, Houston, TX), The hSyn-EGFP plasmid was then modified in-lab by replacing EGFP with mCherry (originally sourced from Addgene #44361). This hSyn-mCherry plasmid was also packaged using a commercial service (Vigene Biosciences, Rockville, MD).

### Focused Ultrasound-mediated Blood-Brain Barrier Opening and Gene Delivery

Mice were anesthetized with 2% isoflurane in air, with the hair on the head shaved with a trimmer and their tail vein cannulated using a 30-gauge needle connected to PE10 tubing. The cannula was flushed with 10 units (U)/ml of heparin in sterile saline (0.9% NaCl) and attached to the mouse tail using tissue glue (3M Vetbond). The mouse was then mounted on a stereotactic platform using ear bars, bite bar and nose cone, all connected to an RK50 (FUS Instruments) system. After disinfecting the site using three alternating scrubs of chlorhexidine scrub and chlorhexidine solution, an incision across the midline scalp was vertically made to expose the skull. Using a guide pointer, bregma-lambda markers were then registered in the RK50 software. The ultrasound parameters used were derived from a published protocol and were 1.5 MHz, 1% duty cycle, and 1 Hz pulse repetition frequency for 120 pulses. Accounting for skull attenuation (18%)^26^, the ultrasound pressure of 1.2 MPa was selected to maximize safety of delivery based on our previous study^27^ and of our preliminary data in the laboratory. Subsequently, AAVs were injected via tail vein into the mice. Within two minutes following injection, the mice were also injected via tail vein with 1.5 × 10^6^ DEFINITY microbubbles (Lantheus Medical Imaging) for a single FUS site, which is what has been used in previous studies^27^. Within thirty seconds of the microbubble injection, the mice were insonated by the transducer coupled to the top of the shaven head via Aquasonic gel (Parker Labs). After insonation is complete, the mice were then placed in the home cage for recovery and following two weeks of gene expression, were euthanized with tissues extracted afterwards.

### Intraparenchymal Injection

After mounting on a stereotaxic frame (Kopf), intraparenchymal co-injections of AAV8 and AAV9.retro were performed using a microliter syringe equipped with a 34-gauge beveled needle (Hamilton) that is installed to a motorized pump (World Precision Instruments). Each AAV was injected unilaterally at a dose of 4×10^8^ viral genomes per gram of body weight to the targeted sites in the nucleus accumbens (AP +1.18 mm, ML +1.25 mm, DV +3.9 mm) and basal pontine nucleus (AP -4 mm, ML +0.4 mm, DV -5.5 mm) infused at a rate of 200 nL/min, and the needle was kept in place for 5 min before removing it from the injection site. The mice were then returned to their home cages for two weeks followed by euthanasia and tissue extraction.

### Histology and Tissue Processing

After cardiac perfusion and tissue extraction, brains and selected peripheral organs (liver, kidneys and spleen) were post-fixed overnight in 10% neutral buffered formalin (NBF). Brains were sectioned coronally for analysis of targeting the nucleus accumbens and sagitally for analysis of targeting the basal pontine nucleus at 40-μm thickness using a cryotome (ref, cryotome). Peripheral organs were sectioned at 100-μm thickness using a vibratome (Leica VT1200S). For analysis of transduction in specific cell types, immunohistochemistry was performed on selected brain sections. Sections were first incubated in a blocking solution of 0.2% Triton X-100 and 10% normal goat serum in 1X phosphate-buffered saline (PBS) for 1 hour followed by overnight incubation with primary antibodies in blocking solution at 4°C. The primary antibodies used were anti-GFAP Alexa Fluor 405-conjugated antibody (1:200 dilution, GFAP Antibody by Novus Biological, stock number: NBP1-05197AF405), and anti-Iba1 Alexa Fluor 405-conjugated antibody (1:200 dilution, Iba1 Antibody by Novus Biological, stock number: NBP1-75760AF405). For oligodendrocyte staining, sections were immunostained with rabbit anti-Olig2 antibody (1:200 dilution, Abcam, stock number: 109186) and Alexa Fluor 647-conjugated goat anti-rabbit IgG antibody (1:200 dilution, Invitrogen, stock number: A21244). The sections were then washed thrice and mounted on glass slides. All sections were then mounted on glass slides with or without DAPI and imaged using a Keyence (Osaka, Japan) BZ-X810 fluorescence microscope under a 20x objective. Brain sections that had transduction at the targeted site and/or exhibited transduction outside the targeted site were selected for analysis and counting.

## Statistical analysis

A two-tailed *t* test was used when comparing the means of two groups. For groups with non-normally distributed data, we used a two-tailed Mann-Whitney test for unpaired datasets. Statistical significance was set at p < 0.05, and software (Prism 9) was used for statistical analysis..

## Data availability

The authors declare that all data supporting the results in this study are available within the paper, its Supplementary Information and its Source Data file. Microscopy images are available from the corresponding author upon reasonable request owing to their large size and numbers.

## ACKNOWLEDGEMENTS

The work described in this study was supported by a grant from the G. Harold and Leila Y. Mathers Charitable Foundation to J.O.S, and by the NSF GRFP award to M.H. Related work in the laboratory was also supported by The Welch Foundation grant to J.O.S. (C-2048-20200401).

## AUTHOR CONTRIBUTIONS

J.O.S. and M.H. conceptualized the study, J.O.S. and M.H. wrote the manuscript J.O.S. and M.H. conceived and planned the research. M.H. and J.O.S. designed the experiments and wrote the paper, with input from the other author. M.H. performed and participated in all experiments described in the study. S.N. performed and participated in intraparenchymal injections and analysis of histology. J.O.S. supervised the study.

## Competing interests

J.O.S. is a coinventor on the pending patent application describing AAV.FUS.3. (US20230047753A1). J.O.S. is a cofounder of Imprint Bio Inc.

## Notes

### Competing Interest Statement

J.O.S. is a coinventor on the pending patent applicationdescribing AAV.FUS.3. (US20230047753A1). J.O.S. is a co-founder of Imprint Bio Inc.

