## Supplementary figures for "Site-Specific Noninvasive Delivery of Retrograde Viral Vectors to the Brain"

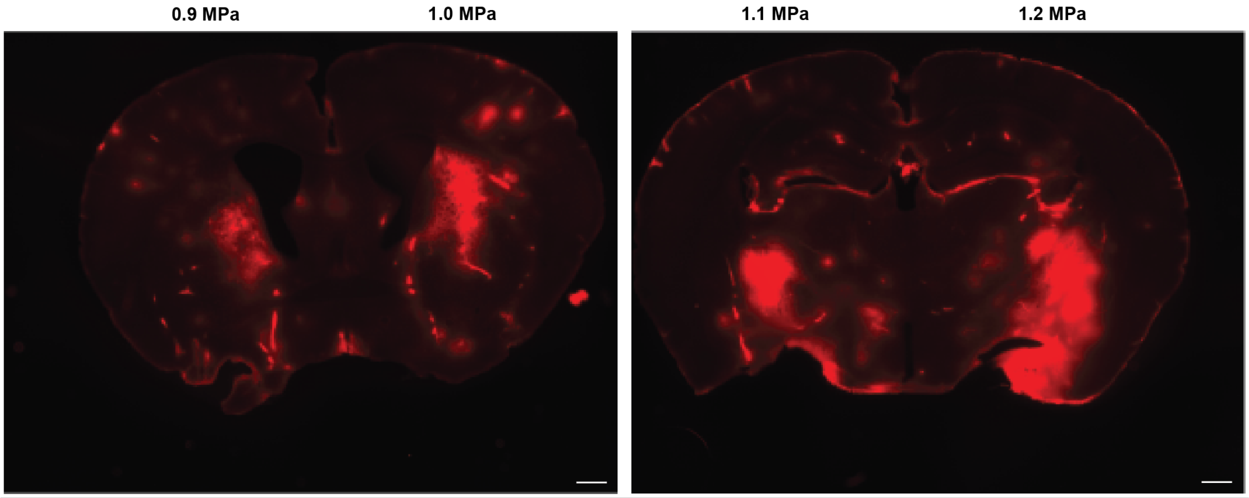


**Supplementary Figure S1: Pilot study for Pressure Optimization for Safe and Effective FUS-BBBO**
Through intravenous injection of Evans Blue Dye (EBD), we tested peak pressures on our FUS system between 0.9 MPa and 1.2 MPa at 1.5 MHz frequency, with 1 Hz pulse repetition frequency for 120 pulses. We observed increased BBB opening with the increase in peak pressure without the presence of hemorrhage and tissue damage, with FUS-BBBO at 1.2 MPa showing both safe and effective extravasation of EBD into the targeted site. We used relatively few animals for this optimization due to the large amount of prior data in our and other groups on the FUS-BBBO. The pressures reported follow the instrument manufacturer's calibration. Scale bars are 500 μm.

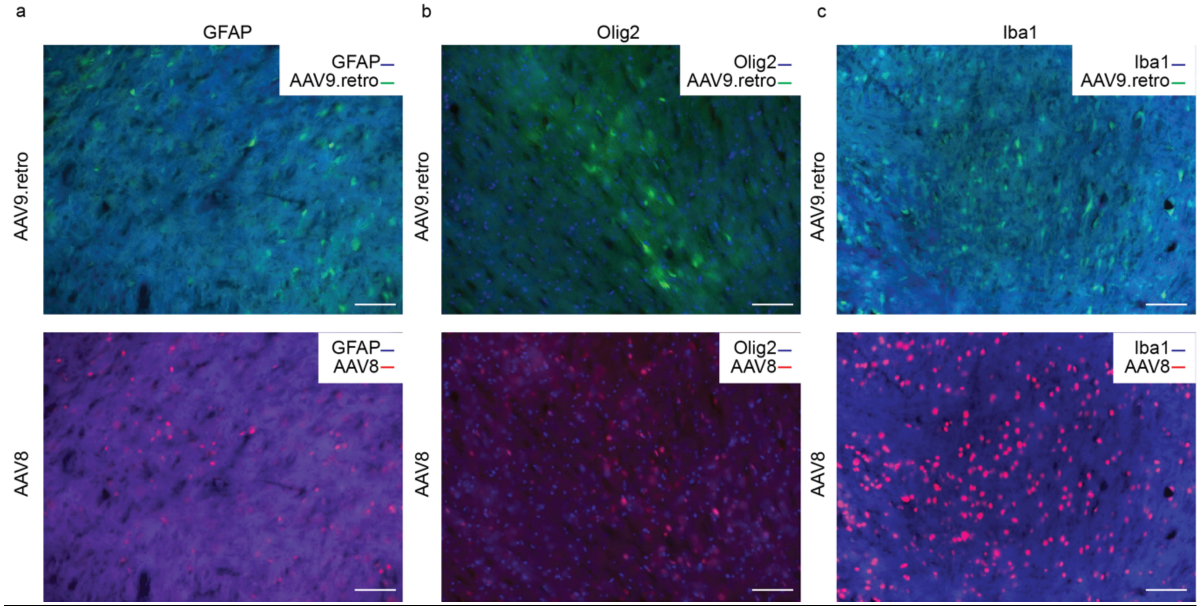


**Supplementary Figure S2: Transduction of non-neuronal cell types by AAV8 and AVV9.retro.**
Since we used a neuron-specific human synapsin promoter to drive expression of mCherry and GFP, and observed no substantial transduction of **a)** astrocytes, **b)** oligodendrocytes and **c)** microglia/macrophages in the brain by either AAV9.retro or AAV8. Representative images were obtained from mice (n=6) co-injected intravenously with AAV8 (red, mCherry) and AAV9.retro (green, EGFP) at $10^{10}$ viral particles per gram of body weight. Sections were imaged on a fluorescence microscope with a 20x objective counterstained for glial cells (GFAP, blue), oligodendrocytes (Olig2, blue) and microglia/macrophages (Iba1, blue). Scale bars are 100 µm.

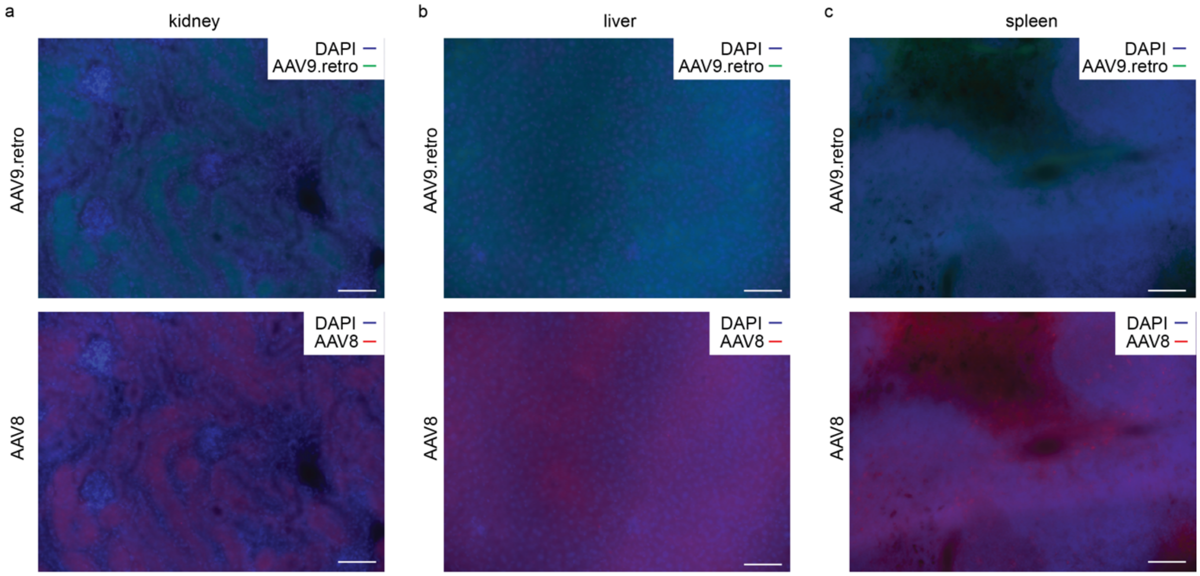


**Supplementary Figure S3: Transduction of viral vectors in peripheral tissues.**
No detectable transduction was observed in the **a)** kidneys, **b)** liver and **c)** spleen. Representative images were obtained from mice (n=8) co-injected intravenously with AAV8 (red, mCherry) and AAV9.retro (green, EGFP) at $10^{10}$ viral particles per gram of body weight. Sections were imaged on a fluorescence microscope with a 20x objective counterstained with a nuclear stain (DAPI, blue). Scale bars are 100 µm.
